# SLIM-2PF: a miniature two-photon fiberscope for multi-depth neuroimaging in freely moving mice

**DOI:** 10.64898/2026.11.12.766309

**Authors:** Jiahan Tian, Yuehan Liu, Chien-Hsien Lai, Hui Lu, Xingde Li

## Abstract

Miniaturized two-photon (2P) fiberscopes enable subcellular-resolution imaging of neural activity in freely moving rodents. However, integrating multi-depth scanning into such probes without compromising their high resolution and minimal mechanical load on the animal remains challenging. Here we report a subcellular-resolution, lightweight, multi-depth-scanning 2P fiberscope (SLIM-2PF). It combines an electrowetting lens with high-numerical-aperture distal optics, achieving lateral and axial resolutions of 0.76-0.81 μm and 6.4-6.8 μm, respectively, across an 84-μm depth range, with a field of view of 168 μm in diameter. By controlling the focus off-head through a thin copper-wire bundle, we reduced the probe weight to only 1.6 g while retaining a flexible tether. In freely moving mice expressing GCaMP6m in pyramidal neurons of the somatosensory cortex, we implemented two multi-depth imaging modes on the same probe: dual-plane recording of dendritic and somatic activity at 3 Hz per plane, and cross-sectional imaging of an unwrapped cylindrical surface spanning 60 μm in depth within each frame at up to 4 Hz, to our knowledge the first demonstration of this mode in freely moving animals. These results establish the SLIM-2PF as a lightweight platform for probing how neural activity is organized across cortical depth during natural behavior.

## 1. Introduction

Recent advances in miniaturized two-photon (2P) devices allow the activity of individual neurons to be imaged in freely moving rodents [1-5], bringing the optical sectioning, deep penetration, and subcellular resolution of 2P microscopy [6-9] to the study of neural circuit dynamics during naturalistic behavior. These head-mounted systems fall into three main architectures. In the first, miniscopes based on a microelectromechanical systems (MEMS) scanner incorporate large-aperture optics on the head, delivering a large field of view (FOV) and high resolution. However, accommodating these optics together with a separate fluorescence-collection path makes the headpiece relatively heavy and bulky [2, 3, 10, 11]. In the second, fiber-bundle endoscopes instead place the scanner at the proximal end, leaving the head end free of any scanner. Yet their discrete core-to-core sampling caps the achievable lateral resolution, and the stiffness of the bundle can restrict the animal’s movement [4, 12, 13].

In the third, fiber-scanning fiberscopes combine a lightweight headpiece with high spatial resolution. They deliver the excitation light through a fiber cantilever whose tip is resonantly scanned in front of a miniature objective [1]. Because excitation and fluorescence share a single double-clad fiber (DCF), no separate collection path is needed on the device, keeping the head end compact [14, 15]. This architecture has enabled neuroimaging in freely moving mice [5, 16], but only within a single focal plane. Cortical activity is organized in three dimensions: different cell types and computations are segregated across cortical layers, and even functionally coupled elements, such as the dendrites and soma of a single neuron, can lie tens of micrometers apart along the depth axis [17-19]. Studying how activity is distributed across depth, and how signals at different depths relate, therefore requires imaging multiple selectable planes during behavior. Optical focus tuning has recently enabled depth-selectable imaging on this platform, but only in head-fixed mice, owing to the bulky tuning driver, and at limited spatial resolution [20]. Realizing multi-depth imaging during free behavior thus calls for integrating this capability compactly into the already space-constrained headpiece, without sacrificing the high resolution, light weight, and flexible tether that make this platform suited to freely moving animals.

Here we address these challenges with a subcellular-resolution, lightweight, multi-depth-scanning 2P fiberscope (SLIM-2PF), built around an electrowetting lens (EWL) and high-numerical-aperture (NA) distal optics. Building on the fiber-scanning platform, we placed the EWL driver off-head and delivered its drive voltage to the probe through a thin copper-wire bundle, reducing the probe weight to 1.6 g, about one-third lighter than the lightest previously reported multi-depth-scanning miniature 2P microscope (Table S1). The high-NA distal optics provide an average resolution of 0.78 μm laterally and 6.6 μm axially, 20% and 34% finer than the best values previously reported for multi-depth-scanning systems in freely moving mice (Table S1). The probe images a 168-μm-diameter FOV at up to 6 Hz per plane, with an 84-μm depth range. In freely moving mice expressing the genetically encoded calcium indicator GCaMP6m in cortical pyramidal neurons, we implemented two multi-depth imaging modes on the same probe: dual-plane recording of dendritic and somatic activity at 3 Hz per plane, and cross-sectional imaging, in which a circular lateral scan synchronized with a continuous focus ramp samples a cylindrical surface that is unwrapped into a depth-spanning image, covering 60 μm in depth in each frame at up to 4 Hz. To our knowledge, this is the first demonstration of cross-sectional functional imaging in freely moving animals. The high resolution and the focus tuning underlying both imaging modes are validated with fluorescent microspheres. Furthermore, we demonstrate volumetric imaging in head-fixed mice and both modes during natural behavior.

## 2. Methods

### 2.1 Probe Design and Fabrication

The SLIM-2PF integrates a resonant fiber scanner, an EWL for focus tuning, and high-NA distal optics into a head-mounted probe (Fig. 1(a)). The rigid distal section measures ∼10 mm in outer diameter and ∼33 mm in length, and the fully assembled probe, excluding the probe holder, weighs 1.6 g (Fig. 1(b)). Excitation light is delivered through the single-mode core of a DCF, whose distal cantilever is driven into resonant spiral scanning by a piezoelectric tube (PZT). The distal end of the DCF is fusion-spliced to a short (∼0.8 mm) silica spacer, onto which a gradient-index lens (GRIN Lens1; GRINTECH IFRL-035-005-50) is bonded. The beam expands freely within the spacer, whose length sets the NA pre-amplification, raising the beam NA from ∼0.12 at the fiber core to ∼0.3 at the output of the composite cantilever. This allows tight focusing with a low-magnification distal objective, enlarging the field of view and imaging throughput without lengthening the cantilever or reducing the scanning speed [21].

**Fig. 1.**
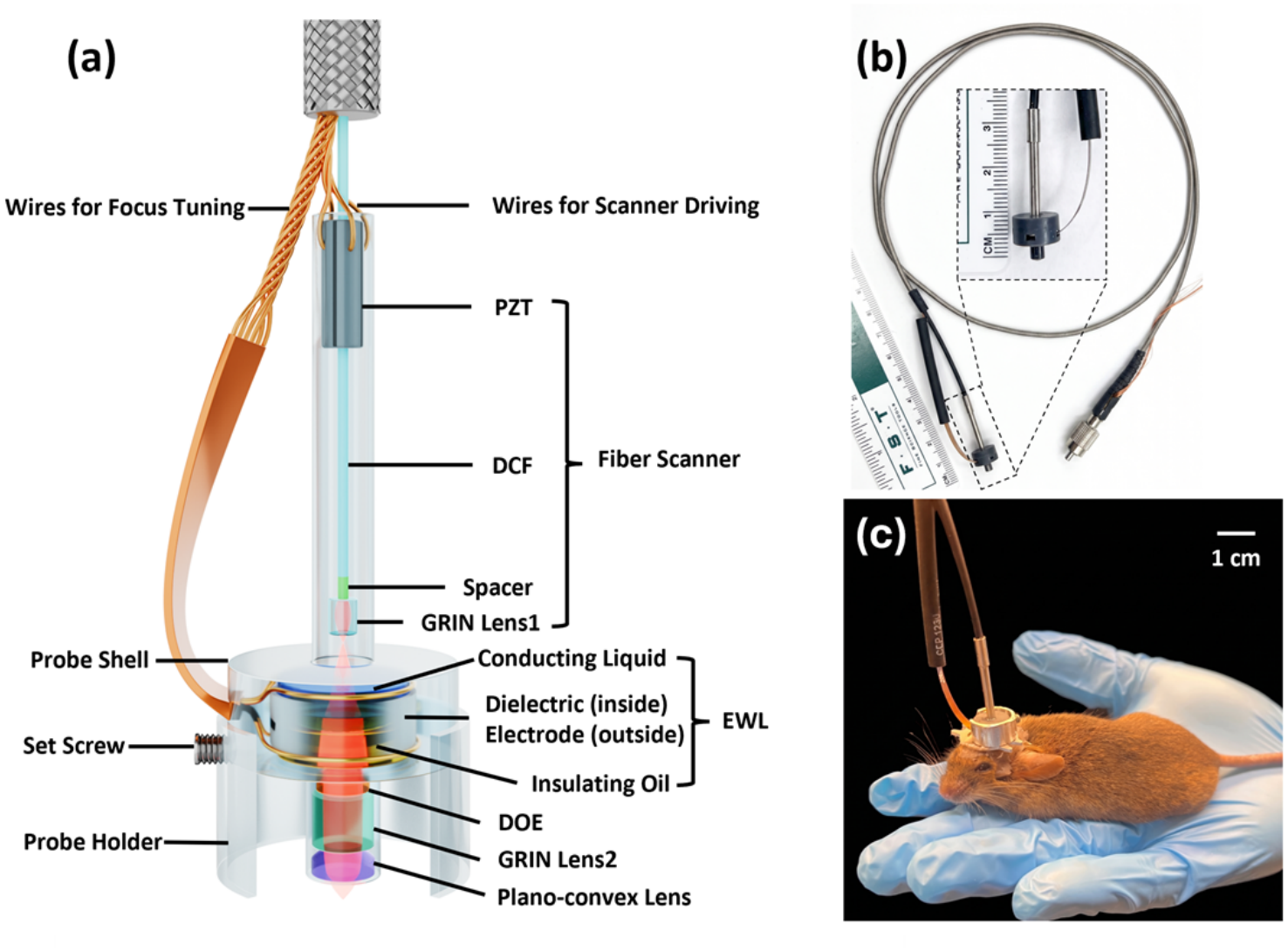
Design and head mounting of the SLIM-2PF. (a) Schematic of the probe. A piezoelectric tube (PZT) resonantly scans a double-clad fiber (DCF), whose distal end is bonded through a short silica spacer to a gradient-index lens (GRIN Lens1) that pre-amplifies the NA. The beam then passes through the electrowetting lens (EWL) for axial focus tuning, a diffractive optical element (DOE) for aberration correction, and a distal objective formed by a second GRIN lens (GRIN Lens2) and a plano-convex lens, which raises the NA at the sample. Ten wires in the tether connect the PZT and the EWL to their off-head drivers. The assembly is housed in a probe shell and secured in a probe holder with a set screw. (b) Assembled probe (1.6 g) with its flexible tether; ruler for scale. (c) Probe mounted on a freely moving mouse. Scale bar in (c), 1 cm.

The beam then passes through the EWL (Corning A-25H0-D0), which tunes the axial focus without moving parts. As shown in Fig. 2(a), the lens contains two immiscible liquids, an upper conducting liquid and a lower insulating oil, whose curved interface forms the optical surface. These two liquids are sealed by a glass window and bounded by a dielectric-coated sidewall with an outer electrode. Applying a voltage between the conducting liquid and the electrode, across the thin dielectric layer, changes the wettability of the liquid on the sidewall (the electrowetting effect) and thereby reshapes the curvature of the oil-liquid interface. Increasing the drive voltage from 34 to 64 V progressively bends this interface from a diverging to a converging surface (Fig. 2(a)), which continuously shortens the focal length and shifts the imaging plane in depth. To keep the probe light, the EWL driver (Corning USB-M Universal V2) is placed off-head and connected to the lens through six soldered copper wires for voltage and temperature control, while the PZT is driven through two wire pairs. All ten wires (each 150 μm in diameter and 0.8 m long) are bundled alongside the DCF in the tether, whose overall diameter, including the protective jacket, is 2 mm.

**Fig. 2.**
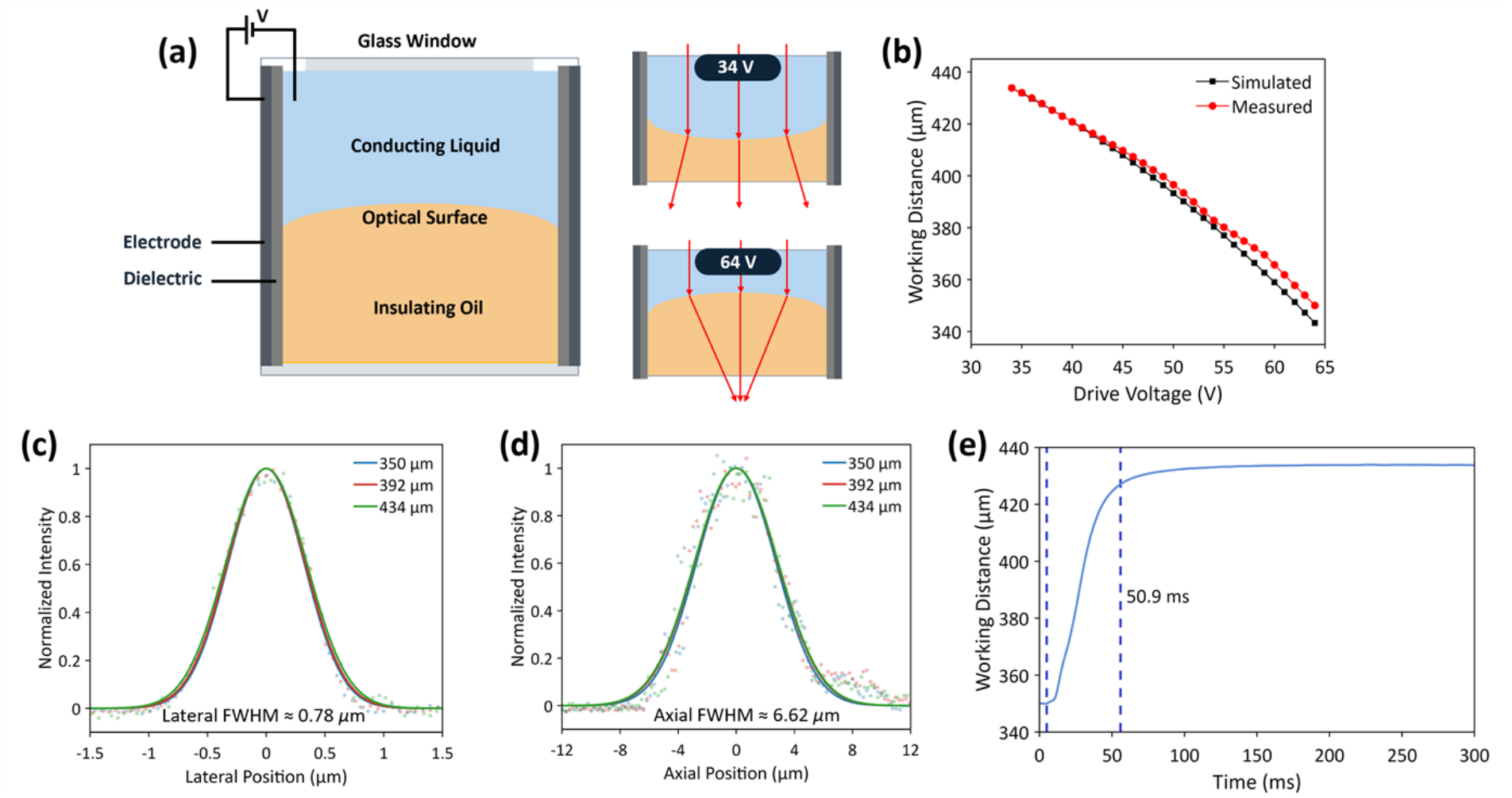
Principle and characterization of multi-depth scanning. (a) Schematic of the EWL (left) and ray-tracing diagrams of the distal optics at drive voltages of 34 and 64 V (right), showing voltage-controlled tuning of the working distance (WD). (b) Measured and Zemax-simulated WD versus drive voltage. (c) Lateral and (d) axial intensity profiles of 200-nm-diameter fluorescent microspheres in water at WD = 350, 392, and 434 μm, with Gaussian fits. Lateral FWHM: 0.76, 0.78, and 0.81 μm; axial FWHM: 6.43, 6.68, and 6.76 μm, respectively. (e) Step response of the WD for a focus switch from 350 to 434 μm; the WD reaches 428 μm (93% of the step) in 50.9 ms.

**Fig. 3.**
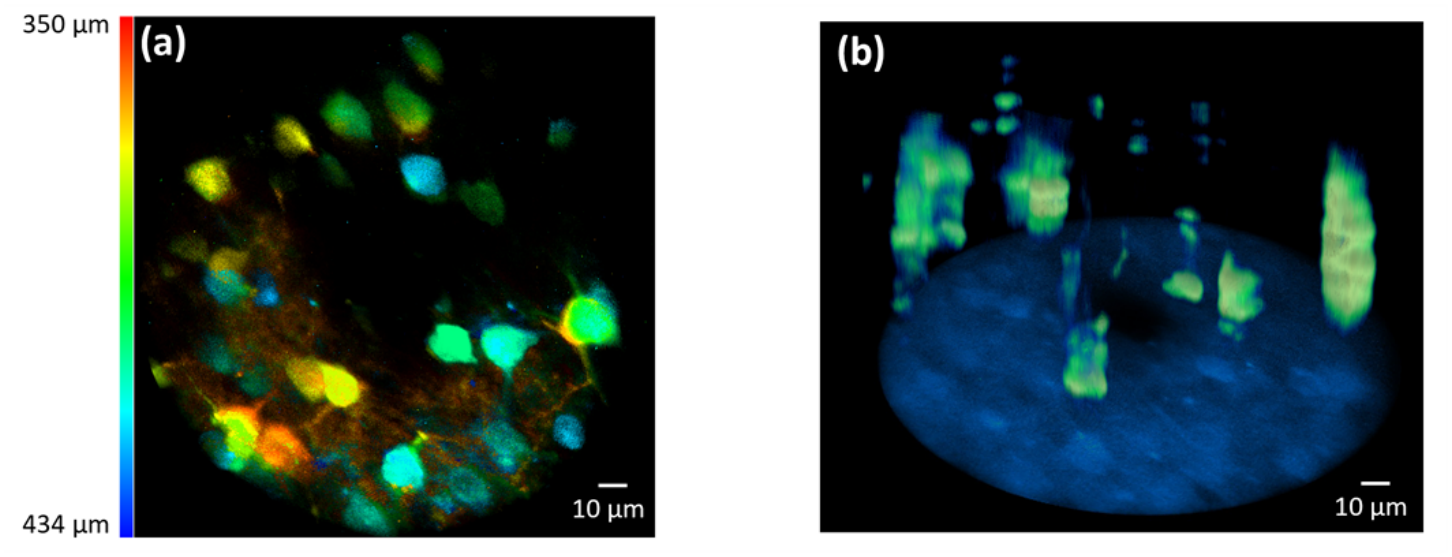
Volumetric imaging of cortical neurons in a head-fixed mouse. (a) Maximum-intensity projection of a z-stack of neurons expressing GCaMP6m in the somatosensory cortex, color-coded by WD from 350 μm (red) to 434 μm (blue). Scale bar, 10 μm. (b) Three-dimensional reconstruction of the same z-stack, viewed at a tilt to show the axial distribution of somata. Scale bar, 10 μm.

Following the EWL, the distal optics are configured to preserve image quality and resolution across the full tuning range. A customized diffractive optical element (DOE) corrects the aberrations introduced by the miniature optics, and a second GRIN lens (GRIN Lens2; GRINTECH LFRL-200-023-50) together with a distal plano-convex lens (Edmund 65297) raises the effective NA at the sample to ∼0.45, thereby improving the spatial resolution. Two-photon fluorescence excited at the focus is collected back through the large-area inner cladding of the same DCF. The fiber scanner is enclosed in a hypodermic metal tube that serves as its outer housing, while the remaining distal optics (the EWL, DOE, GRIN Lens2, and plano-convex lens) are aligned and secured in a 3D-printed probe shell. For *in vivo* use, the assembled probe is locked into a holder with a set screw, and the holder is fixed to the mouse’s head bar with dental cement (Fig. 1(c)). The SLIM-2PF is coupled to a benchtop 2P system: a femtosecond Ti:sapphire source with dispersion pre-compensation delivers 920-nm excitation to the probe, and 2P fluorescence is detected by a photomultiplier (Fig. S1). The average power at the probe output was ∼38 mW during *in vivo* imaging.

### 2.2 System Characterization

The probe was characterized on the bench and with phantoms prior to *in vivo* use. The working distance (WD) as a function of EWL drive voltage was measured by imaging 200-nm-diameter fluorescent microspheres mounted on a motorized linear stage (Newport LTA-HS) in water. At each drive voltage, the stage position of maximum 2P signal was recorded as the WD. The measured WD was compared with a Zemax ray-tracing model of the distal optics. Over a drive range of 34-64 V, the measured WD tuned continuously from 350 to 434 μm, in close agreement with the simulated range of 343-434 μm (Fig. 2(b)), giving a multi-depth scanning range of 84 μm.

Lateral and axial resolutions were measured by imaging 200-nm-diameter fluorescent microspheres and fitting a Gaussian to the lateral and axial profiles of an individual bead to extract the full width at half maximum (FWHM). Measurements were repeated at three working distances spanning the tuning range (350, 392, and 434 μm). The lateral FWHM was 0.76, 0.78, and 0.81 μm, and the axial FWHM was 6.43, 6.68, and 6.76 μm at these three depths, respectively (Figs. 2(c) and 2(d)), indicating subcellular resolution that is well maintained across the full depth-scanning range. The lateral FOV, calibrated by translating a single fluorescent microsphere from one edge of the field to the opposite edge with a precision translation stage, was ∼168 μm in diameter.

The temporal response of focus switching was measured by passing the probe output through a collimator and a pinhole onto a photodetector. Because the collimation of the output beam, and hence the pinhole-transmitted intensity, varies with the WD, the transmitted intensity provides a direct temporal readout of the focal position, which was converted to WD using the calibration in Fig. 2(b). We measured the largest step, from 350 to 434 μm, which represents the worst case; smaller steps are expected to settle faster. The WD reached 428 μm, 93% of the full step, in 50.9 ms (Fig. 2(e)). Because the remaining ∼6 μm is smaller than the axial FWHM at this depth (6.76 μm), we take 50.9 ms as the effective settling time.

### 2.3 Multi-depth Imaging Modes

Because the EWL sets the imaging depth electrically and settles within tens of milliseconds, the axial focus can be reconfigured on the fly and synchronized with the lateral fiber scan. We exploit this to implement two multi-depth imaging modes on the same probe without any change to the optical hardware: dual-plane imaging and cross-sectional imaging. In both modes, the DCF is driven by the PZT to sample the field, and 2P fluorescence is collected continuously through the inner cladding; the modes differ only in the fiber-scan trajectory and in how the EWL drive waveform is synchronized with it.

#### Dual-plane imaging

The fiber is driven in the resonant spiral used for conventional en-face imaging, which acquires a full-FOV plane in ∼110 ms followed by a ∼57-ms fly-back, giving a frame period of ∼167 ms. To image two planes alternately, the EWL drive voltage is toggled at each frame boundary between two preset levels corresponding to WD_1_and WD_2_, so that consecutive frames sample the shallow and deep planes in turn. The focus switch is completed within the fly-back: even the full 84-μm step settles in 50.9 ms (Fig. 2(e)), and the smaller inter-plane steps used here are expected to settle faster, so alternating the focus adds no dead time beyond the fly-back. Each plane is therefore sampled at 3 Hz (6 Hz overall). The interleaved frames are demultiplexed by focus state into two co-registered image series, offset in time by one frame period (∼167 ms), which is short compared with the decay of GCaMP6m transients. Because both planes are imaged through the same distal optics over a common lateral field, structures at different depths, such as a soma and its overlying dendrites, can be imaged and their activity correlated within a single FOV (Fig. 4(a)).

**Fig. 4.**
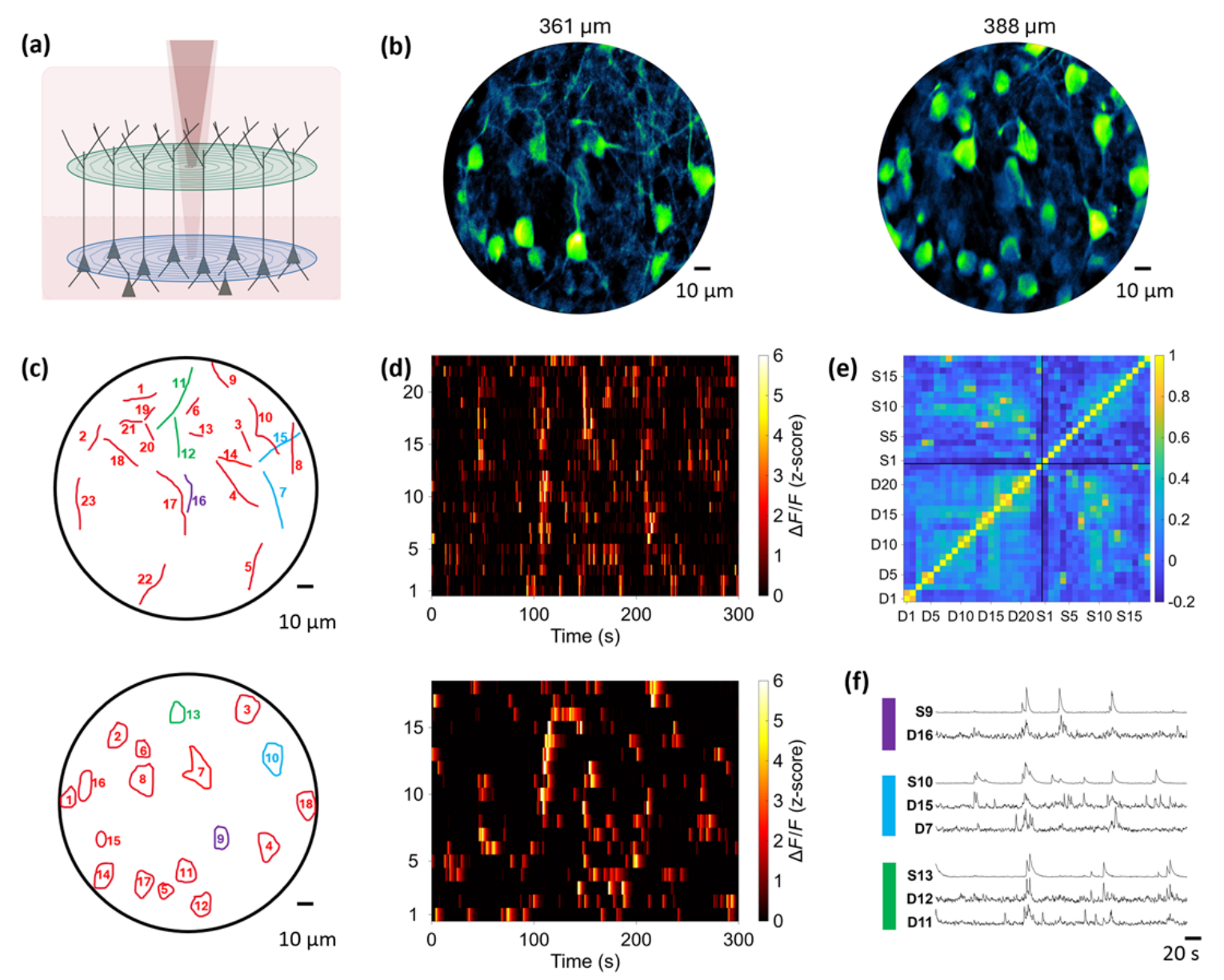
Dual-plane calcium imaging of dendrites and somata in a freely moving mouse (see Visualization 1). (a) Schematic of dual-plane acquisition: the EWL alternates the focus between two axial planes on successive frames, sampling a superficial dendritic plane and a deeper somatic plane. (b) Maximum-intensity projections of the two planes acquired *in vivo* at WDs of 361 μm (dendritic plane) and 388 μm (somatic plane) in mouse cortex expressing GCaMP6m; the imaging planes are estimated to lie within cortical layer 2/3 (L2/3). Scale bar, 10 μm. (c) ROI masks for the dendritic plane (top, 23 putative dendrites) and somatic plane (bottom, 18 somata). (d) Raster plots of z-scored Δ*F*/*F* for all ROIs in the dendritic (top) and somatic (bottom) planes over a 300-s recording. (e) Pearson correlation matrix among all 41 ROIs (D1–D23, dendrites; S1–S18, somata), computed from unnormalized Δ*F*/*F* traces; black lines separate the dendritic and somatic blocks. (f) Δ*F*/*F* traces of three spatially adjacent dendrite–soma groups with correlated activity. Trace colors match the ROI groups in (c) and (e). Normalized per ROI for display. Scale bar, 20 s.

#### Cross-sectional imaging

To sample activity directly along depth, the fiber is driven along a constant-radius circular trajectory, with the two scan axes driven sinusoidally in quadrature at equal amplitude, while the EWL voltage is ramped continuously in synchrony with the circular scan. Within one frame, the fast circular scan repeats many times while the focus is swept once, so the beam traces the lateral surface of a cylinder: the azimuthal position forms the fast lateral axis, and the focal position forms the slow axial axis. Unwrapping this cylindrical surface yields an image spanning the scan circumference (π × 168 μm ≈ 528 μm) and the depth sweep. We refer to this unwrapped cylindrical surface as a cross-section because each frame spans depth. Unlike a conventional planar x–z section, however, it follows a curved lateral path. Because the voltage is swept without dwelling at discrete planes, the axial position of each turn is assigned from a time–depth calibration: the same ramp waveform was applied while recording the pinhole-transmitted intensity with the setup described in Section 2.2, and the trace was converted to WD using the calibration in Fig. 2(b). The ramp rate was chosen to balance axial coverage against frame rate, giving a 60-μm depth sweep per frame at up to 4 Hz. Within one 250-ms frame, the circular scan repeats ∼570 times, so successive turns are separated by ∼0.1 μm in depth, far finer than the 6.6-μm axial FWHM; the cylindrical surface is therefore sampled continuously in depth. To verify that this mode preserves axial resolution and samples depth accurately, we translated a fluorescent microsphere through the sweep with a precision stage: the axial FWHM remained 6.5–6.7 μm at all positions, consistent with the static measurement (Fig. 2(d)), and the measured bead positions closely followed the stage displacements (Fig. S2). Unlike an en-face plane, each frame spans depth, so neurons at different depths are captured within the same 250-ms frame rather than in sequentially acquired z-planes (Fig. 5(a)).

**Fig. 5.**
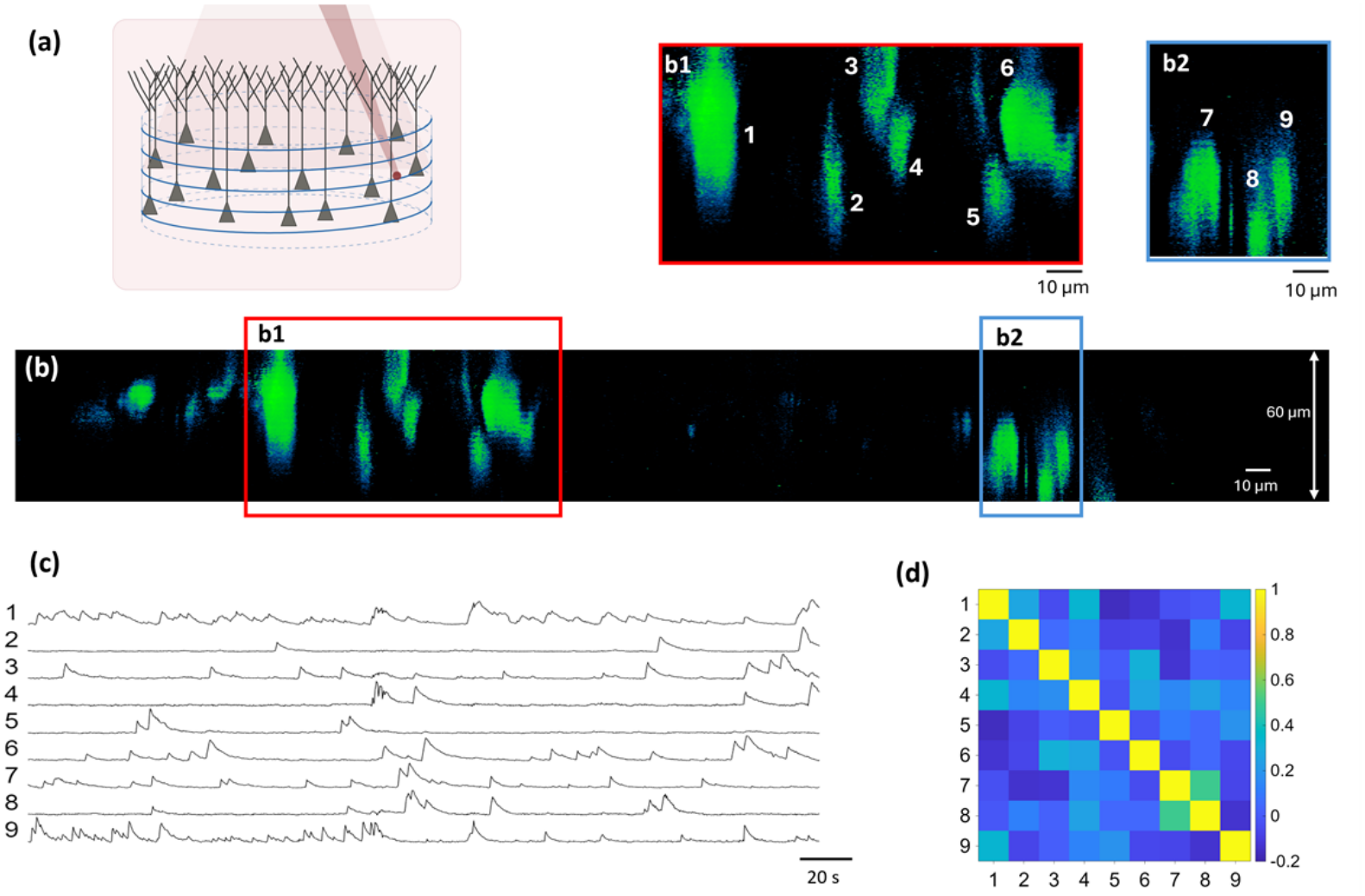
Cross-sectional calcium imaging in a freely moving mouse (see Visualization 2). (a) Schematic of cross-sectional acquisition: the fiber scanner traces a constant-radius circular trajectory while the EWL voltage is ramped continuously over a 60-μm depth range, so that the beam sweeps the lateral surface of a cylinder at 4 Hz. (b) Representative cross-sectional image acquired *in vivo* in the somatosensory cortex expressing GCaMP6m, obtained by unwrapping the cylindrical surface; the horizontal axis spans the scan circumference (528 μm) and the vertical axis the depth sweep (60 μm). The regions in the red (b1) and blue (b2) boxes are shown enlarged above, resolving individual neuronal structures (labeled 1-9) at different depths. Scale bars, 10 μm. (c) Δ*F*/*F* traces of the nine labeled structures, normalized per structure for display. Scale bar, 20 s. (d) Pearson correlation matrix of Δ*F*/*F* among the nine structures, computed from unnormalized traces.

### 2.4 Animal Preparation

All animal procedures were approved by the Institutional Animal Care and Use Committee of Johns Hopkins University and were carried out in accordance with the National Institutes of Health Guide for the Care and Use of Laboratory Animals. Two male Camk2-Cre mice (Jax, #005359) were used: mouse 1 for volumetric imaging under head fixation (Section 3.1), and mouse 2 for dual-plane and cross-sectional imaging during free movement (Sections 3.2 and 3.3). Mice were anesthetized with 2% isoflurane in O_2_ and fixed on a stereotaxic platform. A 4-mm-diameter craniotomy was made over the somatosensory cortex, and AAV/DJ-flex-GCaMP6m (Neuroconnectivity Core, Baylor College of Medicine) was injected through a glass micropipette (TIP10LT, World Precision Instruments) with a microinjector (Nanoject II, Drummond), restricting GCaMP6m expression to CaMKII-positive pyramidal neurons. The exposed cortex was covered with a 100-μm-thick glass coverslip sealed to the skull with tissue adhesive (3M Vetbond), and a custom titanium head bar was attached with dental cement for mounting the probe holder. Expression was verified with a benchtop 2P microscope 3–4 weeks after surgery, before imaging with the SLIM-2PF.

## 3. Results

### 3.1 Volumetric Imaging in Head-fixed Mice

We first verified that the multi-depth scanning range translates into volumetric imaging in tissue. In an awake, head-fixed mouse expressing GCaMP6m in the somatosensory cortex, a z-stack was acquired by advancing the WD in 2.8-μm steps, with the drive voltage for each step set from a polynomial fit to the voltage–WD calibration (Fig. 2(b)). The frames were stacked to reconstruct the volume. Because the 2.8-μm step is less than half the 6.6-μm axial FWHM, the volume is sampled continuously in depth.

Figure 3(a) shows the depth-color-coded maximum-intensity projection of the stack. Neurons at different depths appear in different colors and remain individually distinguishable, and both somata and dendrites are resolved throughout the stack, confirming that focus tuning remains effective in scattering tissue. The three-dimensional reconstruction (Fig. 3(b)) resolves ∼20 somata, distributed across the 84-μm depth range, estimated to lie within layer 2/3. This depth discrimination allows the EWL to be set to a chosen drive level to target neurons at a specific depth, as exploited in the dual-plane mode below.

### 3.2 Dual-plane Imaging of Dendritic and Somatic Activity

We next applied the dual-plane mode to the somatosensory cortex of a freely moving mouse. The EWL was toggled between WDs of 361 and 388 μm on successive frames, giving an inter-plane separation of 27 μm; both planes are estimated to lie within cortical layer 2/3. The shallower plane captured a dense field of neuropil and dendritic processes, while the deeper plane resolved individual somata (Fig. 4(b)). Each plane was updated at 3 Hz for a 300-s recording without any adjustment to the probe, while the mouse moved freely in the arena with no apparent impairment of posture or locomotion (Visualization 1). All frames were motion-corrected with NoRMCorre [22] before segmentation. ROIs were delineated manually on the local correlation image of each plane, yielding 23 dendritic and 18 somatic ROIs (Fig. 4(c)). For each ROI, Δ*F*/*F* was computed as (*F* − *F*_*0*_)/*F*_*0*_, where *F* is the mean fluorescence within the ROI, and *F*_*0*_ is a baseline estimated by a running minimum followed by a running median over a 30-s window. The dendritic ROIs were identified as putative dendrites on the basis of their thin, elongated, and branching morphology and their location above the somatic plane, consistent with the apical dendrites of CaMKII-positive L2/3 pyramidal neurons. Without a structural marker, however, some of these ROIs may also include axonal processes. Calcium transients were observed in both planes throughout the recording; the raster plots and Δ*F*/*F* traces in Figs. 4(d) and 4(f) are normalized per ROI for display clarity. The somatic traces appear smoother than the dendritic traces because each somatic ROI averages the fluorescence over many more pixels than a thin dendritic ROI, which reduces the noise in the extracted trace.

Because the two planes are imaged through the same distal optics over a common lateral field, dendritic and somatic signals can be related directly. We computed the pairwise Pearson correlation between all dendritic and somatic ROIs using the unnormalized Δ*F*/*F* traces (Fig. 4(e)). Several spatially adjacent dendrite-soma pairs showed correlated activity: D12-S13 (r = 0.63) and D11-S13 (r = 0.58); D15-S10 (r = 0.52) and D7-S10 (r = 0.49); and D16-S9 (r = 0.60). Their Δ*F*/*F* traces co-vary closely, with dendritic and somatic transients often occurring within the same acquisition cycle (Fig. 4(f)). The spatial proximity and correlated activity suggest that each group may belong to a single neuron or to cells within a common local circuit. Such relationships cannot be examined with a single-plane fiberscope, which records only one depth at a time.

### 3.3 Cross-sectional Imaging

Finally, we tested the cross-sectional mode in the somatosensory cortex of a freely moving mouse. A single continuous focus ramp synchronized to the circular scan produced one depth-spanning frame at 4 Hz. The unwrapped frames were motion-corrected and processed as described in Section 3.2, with ROIs delineated manually on the mean cross-sectional image. The resulting cross-sections resolve individual neuronal structures at clearly different depths within a single frame (Fig. 5(b)); the enlarged regions (b1, b2) show nine such structures distributed across the 60-μm depth sweep. Because every frame covers a cylindrical surface of 528 × 60 μm^2^, structures that would fall in separate z-planes of a conventional acquisition appear together in one frame. Some of these structures appear axially elongated. This likely reflects several contributions rather than the true extent of a single soma: vertically oriented neuronal processes (axons and dendrites running along the optical axis) that are sampled across the depth sweep; somata that overlap along depth and merge into a single elongated feature; and axial tailing from the finite 6.6-μm axial resolution. Consistent with the first contribution, elongated activity regions attributed to axonal or dendritic processes in the neuropil have been reported at shallow cortical depths with an EWL-based fiber-coupled 2P microscope [13].

The Δ*F*/*F* traces of the nine structures (Fig. 5(c)) show distinct calcium dynamics during free behavior (Visualization 2), with sparse and largely independent activity. Pairwise correlations computed from unnormalized traces (Fig. 5(d)) were close to zero for most pairs, except for the neighboring pair #7-#8 (r = 0.50), located at different depths, whereas the nearby structure #9 was uncorrelated with both. This pattern suggests that the correlated activity of #7 and #8 may reflect a functional relationship rather than a global artifact from animal motion or shared background, which illustrates that the cross-sectional mode can relate the activity of structures at different depths within the same frame.

## 4. Discussion

The key design choice of the SLIM-2PF is where the depth degree of freedom is placed and how it is driven. Rather than translating the optics or the probe itself, the focus is set by the curvature of a liquid–liquid interface inside the EWL, and the EWL driver is placed off-head, so that depth becomes an electrical parameter delivered through a thin copper-wire bundle. This avoids both the moving mass of mechanical focus scanning and a bulky on-head driver, allowing multi-depth scanning to be integrated without compromising the high resolution, light weight, and flexible tether that make fiber-scanning fiberscopes suited to freely moving animals. Multi-plane imaging in freely moving mice has been demonstrated previously with MEMS-based miniscopes [2, 10]. The contribution here is to bring multi-depth scanning, including a mode that captures cross-sectional structure within a single frame, to the fiber-scanning fiberscope at a probe weight of only 1.6 g.

The two modes give the SLIM-2PF a capability that a single en-face plane lacks: relating activity at one depth to activity at another. Dual-plane imaging does this for two discrete, chosen depths, a superficial dendritic (input) plane and a deeper somatic (output) plane, sampled near-simultaneously across a small axial separation, supporting the study of dendritic integration and input-output coupling within a single field while also doubling the structures recorded per acquisition. Cross-sectional imaging extends the same idea to a continuous depth range within one frame: because different depths are sampled within the same frame, a single cross-section shows how activity is distributed along the cortical axis, matching the depth organization of cortical circuits, and can relate activity between structures separated in depth. A longer focus-tuning range would extend this single-frame cross-section continuously across multiple cortical layers.

Because the two modes share identical hardware and differ only in the drive waveform, the probe can be reconfigured on the fly during a single session. A natural workflow is to use the cross-sectional mode as a survey: one continuous ramp reports which depths contain active structures along the sampled cylindrical surface, after which the probe can switch, with no hardware change, to dual-plane recording at the two depths of interest for sustained observation. The same waveform flexibility admits other trajectories on the same probe, such as multi-radius cylindrical surfaces that sample more of the volume, or tilted planes matched to the orientation of a cortical column or a labeled projection, extending the accessible sampling geometries without any change to the optics.

While the present recordings show that the SLIM-2PF does not prevent the mouse from moving freely in the arena (Visualizations 1 and 2), more extensive behavioral profiling would further strengthen future studies using this platform. Several applications follow from combining multi-depth imaging with free behavior. First, because the probe imposes only a light, flexible tether, imaging can be paired with synchronized behavior tracking of locomotion, pose, and task events, and the multi-depth activity can be aligned to these behavioral variables on a common timebase, linking what is recorded across cortical depth to what the animal is doing. Second, resolving somatic activity at multiple depths within the same session makes it possible to relate cross-layer activity to behavior or to targeted stimulation, addressing how signals at different cortical depths co-vary as behavior unfolds. Third, the platform is suited to concrete circuit questions that require both subcellular resolution and natural behavior, for example, how dendritic computation contributes to somatic output during behavior or learning, and, because depth is set electrically and reproducibly rather than by mechanical positioning, the same depth-distributed population can be revisited across days for longitudinal imaging.

## 5. Conclusion

We have developed the SLIM-2PF, a fiber-scanning 2P fiberscope that brings multi-depth scanning to freely moving mice. Placing the EWL driver off-head and delivering its drive voltage through a thin copper-wire bundle turns depth into an electrical parameter rather than a moving mass, so that multi-depth scanning is achieved with a probe weight of only 1.6 g, about one-third lighter than the lightest previously reported multi-depth-scanning miniature 2P microscope, while retaining a flexible tether. Despite its low weight, the probe provides the highest resolution among multi-depth-scanning systems for freely moving mice, with a mean of 0.78 μm laterally and 6.6 μm axially, over a 168-μm-diameter FOV and an 84-μm depth range, with focus switching within tens of milliseconds.

On this single platform, without any change to the optical hardware, we implemented two multi-depth imaging modes in freely moving mice: dual-plane recording, which revealed correlated activity between spatially adjacent dendrites and somata across a 27-μm axial separation at 3 Hz per plane; and cross-sectional imaging, which resolved neuronal structures distributed across a 60-μm depth sweep at 4 Hz, to our knowledge the first demonstration of this mode in freely moving animals. Together, these results establish the SLIM-2PF as a lightweight, high-resolution platform for probing how neural activity is organized across cortical depth during natural behavior.

## Supporting information

Visualization 1: Dual-plane imaging of dendrites and somata in a freely moving mouse

Visualization 2: Cross-sectional imaging in a freely moving mouse

## Data availability

Data underlying the results presented in this paper are not publicly available at this time but may be obtained from the authors upon reasonable request.

## Supplemental document

See Supplement 1 for supporting content.

