## Supplement 1: Table S1 and Figures S1 and S2 for "SLIM-2PF: a miniature two-photon fiberscope for multi-depth neuroimaging in freely moving mice"

### Table S1. Miniature two-photon systems with a focus-tuning capability in freely moving mice

| Device | Weight[α] (g) | Scanning method | Lateral resolution (μm) | Axial resolution (μm) | Field of view[β] (μm) | Focus tuning range (μm) |
|---|---|---|---|---|---|---|
| 2P-FCM[1] | 2.5 | Proximal galvo with fiber bundle | 1.8 | 10 | Ø 240 | 180 |
| FHIRM-TPM 2.0[2] | 4.2 | Distal MEMS | 1.13 | 12.18 | 420 × 420 | 180 |
| MINI2P[3] | 2.4 | Distal MEMS | 1.15 - 1.24 | 12.8 - 17.8 | 420×420 (MINI2P-F)<br>500×500 (MINI2P-L) | 240 |
| Zhao *et al.*[4] | 3.7 | Distal MEMS | 1.47 | 24.64 | 1000 × 788 | 500 |
| UCLA 2P Miniscope[5] | 4 | Distal MEMS | 0.98 | 10.18 | 445 × 380 | 150 |
| **SLIM-2PF (This work)** | **1.6** | **Distal piezoelectric fiber scanner** | **0.76 - 0.81** | **6.43 - 6.76** | **Ø 168** | **84** |

[α]All weights include the focus-tuning module.

[β]Ø = circular FOV diameter.

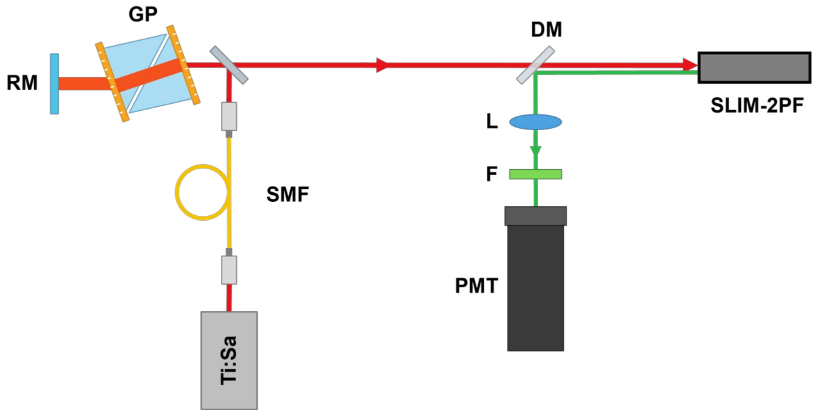


Figure S1. Schematic of 2P fiberscope imaging system. SMF: single-mode fiber; RM: roof mirror; GP: GRISM pair for dispersion compensation; DM: dichroic mirror; L: lens; F: optical filter; PMT: photomultiplier tube.

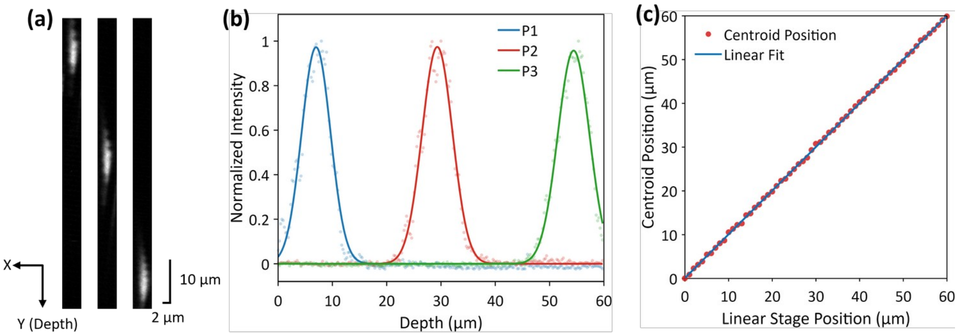


Figure S2. Characterization of cross-sectional imaging with a 200 nm fluorescent microsphere on a precision translation stage. A fluorescent microsphere was imaged in cross-sectional mode while translated along the depth axis with a high-precision stage (LTA-HS, Newport). The PSF centroid at the top edge of the field was taken as the zero reference. (a) Cross-sectional images of the microsphere at three relative positions (7, 30, and 54 µm), all shown on the same scale (scale bars: 10 µm vertical, 2 µm horizontal); the PSF profiles are closely similar across positions. (b) Normalized intensity profiles along the Y (depth) axis with Gaussian fits, yielding axial FWHMs of 6.48 µm (P1), 6.56 µm (P2), and 6.65 µm (P3), consistent with the axial resolution measured on the bench (Fig. 2(d)). (c) PSF centroid position versus commanded stage position, with a linear fit; the measured centroids closely follow the stage displacements, confirming that cross-sectional imaging is spatially uniform and stable across the depth sweep.
